# The circadian clock component BMAL1 regulates SARS-CoV-2 entry and replication in lung epithelial cells

**DOI:** 10.1101/2021.03.20.436163

**Authors:** Xiaodong Zhuang, Senko Tsukuda, Florian Wrensch, Peter AC Wing, Mirjam Schilling, James M Harris, Helene Borrmann, Sophie B Morgan, Jennifer L Cane, Laurent Mailly, Nazia Thakur, Carina Conceicao, Harshmeena Sanghani, Laura Heydmann, Charlotte Bach, Anna Ashton, Steven Walsh, Tiong Kit Tan, Lisa Schimanski, Kuan-Ying A Huang, Catherine Schuster, Koichi Watashi, Timothy SC Hinks, Aarti Jagannath, Sridhar R Vausdevan, Dalan Bailey, Thomas F Baumert, Jane A McKeating

## Abstract

The COVID-19 pandemic, caused by SARS-CoV-2 coronavirus, is a global health issue with unprecedented challenges for public health. SARS-CoV-2 primarily infects cells of the respiratory tract, via Spike glycoprotein binding angiotensin-converting enzyme (ACE2). Circadian rhythms coordinate an organism’s response to its environment and can regulate host susceptibility to virus infection. We demonstrate a circadian regulation of ACE2 in lung epithelial cells and show that silencing BMAL1 or treatment with a synthetic REV-ERB agonist SR9009 reduces ACE2 expression and inhibits SARS-CoV-2 entry. Treating infected cells with SR9009 limits viral replication and secretion of infectious particles, showing that post-entry steps in the viral life cycle are influenced by the circadian system. Transcriptome analysis revealed that Bmal1 silencing induced a wide spectrum of interferon stimulated genes in Calu-3 lung epithelial cells, providing a mechanism for the circadian pathway to dampen SARS-CoV-2 infection. Our study suggests new approaches to understand and improve therapeutic targeting of SARS-CoV-2.

---

Circadian signalling exists in nearly every cell and is primarily controlled by a series of transcription/translation feedback loops. The transcriptional activators BMAL1 and CLOCK regulate thousands of transcripts including their own repressors, REV-ERBα and REV-ERBβ, that provide a negative feedback loop to control gene expression. Human lung diseases frequently show time-of-day variation in symptom severity and respiratory function (Scheiermann et al., 2018) and BMAL1 is recognised to play a key role in regulating pulmonary inflammation (Ince et al., 2019). Influenza A infection of arrhythmic mice, lacking BMAL1, is associated with a higher viral burden in the lung (Edgar et al., 2016) and elevated inflammatory responses (Ehlers et al., 2018; Sengupta et al., 2019). We (Sengupta et al., 2021) and others (Maiese, 2020; Ray and Reddy, 2020) hypothesized a potential role for the circadian clock in COVID-19. To investigate a potential role for circadian pathways in the SARS-CoV-2 life cycle we assessed whether infection was time-dependent in Calu-3 lung epithelial cells. Several approaches have been reported to synchronise the circadian clock in cell culture and serum shocking Calu-3 cells was the optimal method to coordinate Bmal1 promoter activity (**Figure 1A**). Lentiviruses can incorporate exogenous viral glycoproteins and the resulting pseudoparticles (pp) undergo a single cycle of infection that enable the study of receptor-specific internalisation pathways. Infecting synchronized Calu-3 cells at different circadian times (CT) with pseudoparticles expressing the SARS-CoV-2 Spike glycoprotein showed a rhythmic pattern of infection, whereas those bearing the vesicular stomatitis virus (VSV) G glycoprotein infected the cells with comparable efficiency at all time points (**Figure 1B**). Given the reported link between the circadian clock and cell cycle (Farshadi et al., 2020; Masri et al., 2013) that may influence SARS-CoV-2 infection (Bouhaddou et al., 2020; Su et al., 2020), we evaluated the effect of our serum shock protocol on the Calu-3 cell cycle. Serum shock had no significant effect on the Calu-3 cell cycle (**Figure S1A**). These data uncover a potential role for circadian pathways to regulate Calu-3 susceptibility to SARS-CoV-2 uptake.

**Figure 1.**
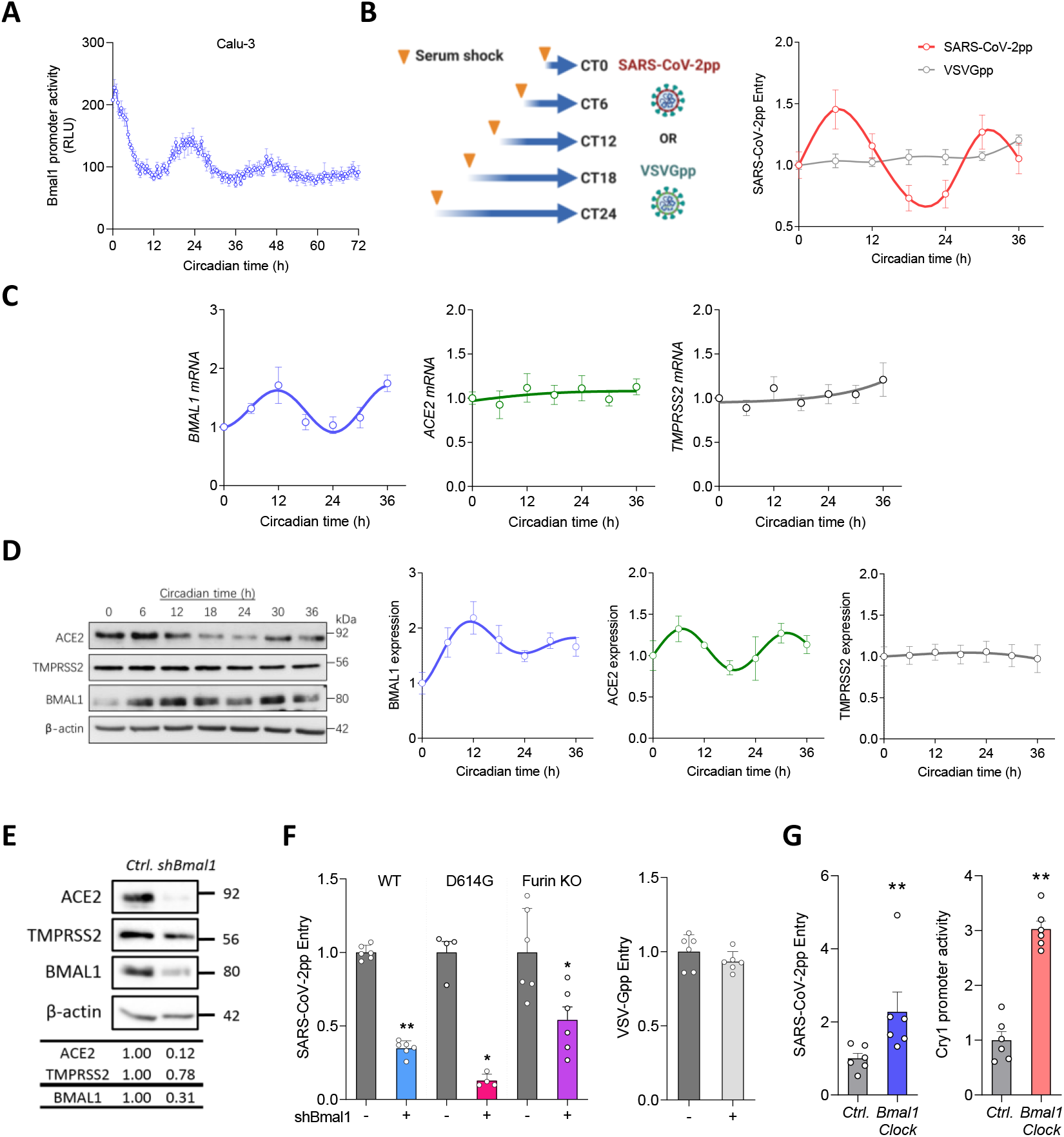
BMAL1 regulation of ACE2 dependent SARS-CoV-2 entry. (**A**) Synchronisation of Calu-3 cells. Calu-3 cells expressing a Bmal1 promoter driven luciferase reporter were synchronised by serum shock (50% FBS) for 1h followed by real-time monitoring of luciferase activity. Data are expressed relative to circadian time 0 (defined by 12h post-serum shock) and are the average of six independent experiments (n = 6, mean ± S.E.M.). (**B**) Circadian infection of SARS-CoV-2pp. Synchronised Calu-3 cells were inoculated with SARS-CoV-2pp or VSV-Gpp for 2h with luciferase activity measured 24h later. Data expressed relative to CT0; mean ± S.E.M., n = 6-12. (**C**) Synchronised Calu-3 cells were assessed for *BMAL1, ACE2* and *TMPRSS2* mRNA by qRT-PCR and expressed relative to CT0; data are the average of four independent experiments. (**D**) Synchronised Calu-3 cells were assessed for ACE2, TMPRSS2 and BMAL1 expression together with housekeeping β-actin by western blotting; data are representative of four experiments. Densitometric analysis quantified ACE2, TMPRSS2 and BMAL1 in individual samples and normalized to their own β-actin loading control. (**E**) Calu-3 cells were transduced with lentivirus encoding shBmal1 or control and ACE2, TMPRSS2, BMAL1 and β-actin protein expression assessed by western blotting. (**F**) Control or shBmal1 treated cells were infected with lentiviral pseudotypes expressing wild type (WT) or mutant (D614G or Furin KO) Spike variants or with VSV-Gpp. Viral entry (luciferase activity) was measured 24h later and data expressed relative to WT (mean ± S.E.M., n = 4-6, Mann–Whitney test). (**G**) Calu-3 cells transfected with Bmal1/Clock expression plasmids were infected with SARS-CoV-2pp or co-transfected with Cry1 promoter luciferase reporter. 48 h post-transfection, cells were lysed and viral entry or promoter activity determined by quantifying luciferase activity. Data are expressed relative to control treatment (mean ± SEM, n = 6, Mann–Whitney test). See also Figure S1 and S2.

Since ACE2 and transmembrane protease serine 2 (TMPRSS2) co-regulate SARS-CoV-2 internalisation (Hoffmann et al., 2020b; Wan et al., 2020) we measured their expression in synchronised Calu-3 cells. Since the classical model of circadian regulation is one of transcriptional control we quantified ACE2 and TMPRSS2 transcripts in synchronised Calu-3 cells and showed no evidence of rhythmic expression, whereas we observed the expected pattern of Bmal1 RNA (**Figure 1C**). However, ACE2 protein expression varied over a circadian cycle (**Figure 1D**), with the peak (CT6) and trough (CT24) of expression associating with SARS-CoV-2pp entry, consistent with a post-transcriptional regulation of ACE2. In contrast, TMPRSS2 expression was similar at all time points sampled (**Figure 1D**). To extend these observations we quantified *Ace2* and *Tmprss2* transcripts in lung and liver harvested from light/dark entrained mice and observed limited evidence for a circadian pattern of expression (**Figure S2**). Immunoblotting of ACE2 in the murine lung resulted in multiple bands of varying molecular weight, compromising the interpretation of these experiments.

To further explore the role of circadian pathways in regulating ACE2 we shRNA silenced Bmal1, the major circadian transcriptional activator, and showed reduced ACE2 expression but a negligible effect on TMPRSS2 in Calu-3 cells (**Figure 1E**). To assess the impact of Bmal1 on SARS-CoV-2pp entry we infected the silenced Calu-3 cells and showed a significant reduction in pp infection, whereas VSV-Gpp infection was unaffected (**Figure 1F**). During the COVID-19 pandemic several Spike variants have emerged; some conferring a fitness advantage to viral entry. D614G, has become prevalent in many countries, consistent with a reported advantage for infecting cells of the upper respiratory tract (Korber et al., 2020). SARS-CoV-2 Spike has a unique furin cleavage site that mediates membrane fusion and deletion of this motif is associated with reduced pathogenesis (Johnson et al., 2021). Importantly, pp containing either Spike variant showed reduced infection of Bmal1 silenced Calu-3 cells (**Figure 1F**). To further explore a regulatory role for Bmal1 in SARS-CoV-2 entry we overexpressed Bmal1 along with Clock in Calu-3 cells and showed a significant increase in both SARS-CoV-2pp entry and promoter activity of the host target Cry1 (**Figure 1G**). In summary, these data show a role for BMAL1 to regulate ACE2 and SARS-CoV-2 entry.

The availability of a synthetic agonist (SR9009) that activates REV-ERB and modulates circadian pathways (Solt et al., 2012; Trump et al., 2013) prompted us to investigate its role in SARS-CoV-2 infection. Treating Calu-3 cells with SR9009 reduced BMAL1 promoter activity and protein expression, with no effect on cell viability (**Figure S3**). SR9009 treatment reduced ACE2 in a dose-dependent manner but had no effect on TMPRSS2 expression (**Figure 2A**). Furthermore, SARS-CoV-2pp infection was significantly reduced by SR9009 treatment in parental Calu-3 cells but not in shBmal1 silenced cells, demonstrating a Bmal1-dependency (**Figure 2B**). In contrast, SR9009 treatment had no effect on VSV-Gpp infection (**Figure 2B**). SR9009 treatment inhibited the infection of pp bearing the D614G or Furin-KO Spike variants (**Figure 2C**). Using an independent pseudoparticle system based on VSV (Hoffmann et al., 2020b), we show that SR9009 treatment of Calu-3 reduced particle infection (**Figure 2D**). To extend our observations to a more physiologically relevant system, we treated differentiated air-liquid interface (ALI) cultures of proximal airway epithelial cells with SR9009 and showed a significant reduction in SARS-CoV-2pp infection and ACE2 expression (**Figure 2E**).

**Figure 2.**
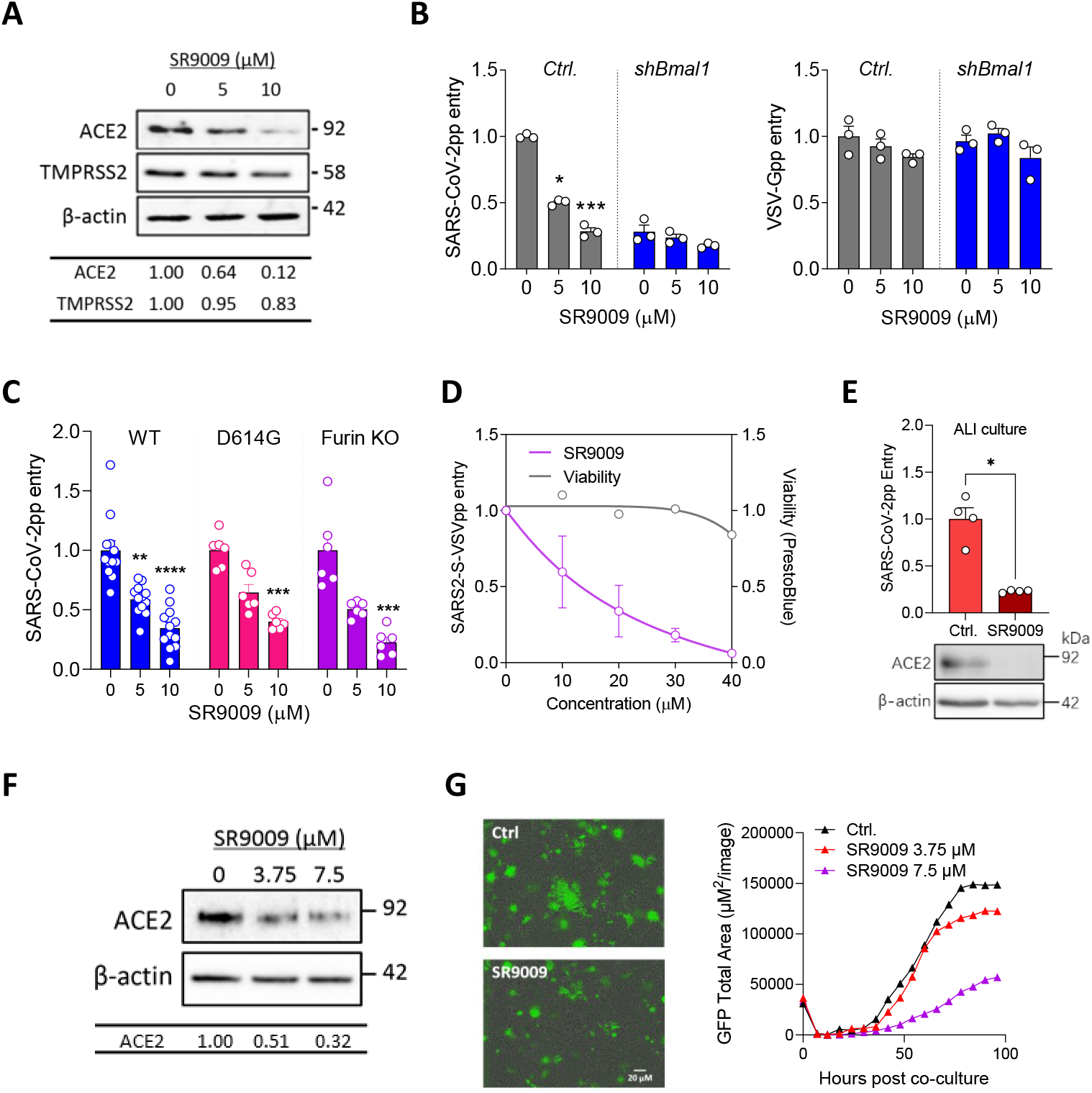
REV-ERB agonist reduces ACE2 dependent SARS-CoV-2 entry and cell-cell fusion. (**A**) Calu-3 cells were treated with SR9009 (5 or 10 µM) for 24h and assessed for ACE2 and TMPRSS2 expression together with housekeeping β-actin by western blotting. (**B**) Control or shBmal1 silenced Calu-3 cells were treated with SR9009 for 24h followed by infection with SARS-CoV-2pp or VSV-Gpp. After 24h viral entry was assessed by measuring luciferase activity and expressed relative to untreated cells (mean ± S.E.M., n = 3, Kruskal–Wallis ANOVA with Dunn’s test). (**C**) Control or SR9009 treated Calu-3 cells were infected with SARS-CoV-2 wild type (WT), D614G or furin mutant pp. Viral entry was expressed relative to untreated cells 24h later (mean ± S.E.M., n = 4-6, Kruskal– Wallis ANOVA with Dunn’s test). (**D**) Control or SR9009 treated Calu-3 cells were infected with SARS2-S-VSVpp and 24h later luciferase activity measured and data expressed relative to untreated cells (mean ± S.E.M., n = 3). Cell viability was determined using PrestoBlue cell viability assay. **(E)** Differentiated air-liquid interface cultures of proximal airway epithelial cells were treated with SR9009 (20 µM) for 24h prior to SARS-CoV-2pp infection. Viral entry was measured as luciferase activity and data presented as mean ± S.E.M of four wells from a single donor relative to untreated control cells (mean ± S.E.M., n = 4, Mann–Whitney test). ACE2 expression together with housekeeping β-actin by western blotting. **(F)** SR9009 treated Huh-7 cells were assessed for ACE2 expression together with the housekeeping gene β-actin. Data are representative of three experiments. Densitometric analysis quantified ACE2 in individual samples and normalized to the β-actin loading control. Representative of three experiments. **(G)** Huh-7 target cells expressing a split rLuc-GFP reporter (8-11) were treated overnight with SR9009 at the indicated concentrations or with DMSO vehicle (Ctrl.) before culturing with Spike expressing HEK-293T cells with the split rLuc-GFP (1-7) and SR9009 added for the indicated times. A representative image of SARS-CoV-2 Spike induced syncytia at 3 days post co-culture is shown (left panel). GFP-positive syncytia were quantified every 4h using an IncuCyte real-time imaging platform (right panel). Five fields of view were obtained per well at 10x magnification and GFP expression quantified by calculating the total GFP area using the IncuCyte analysis software. See also Figure S3.

SARS-CoV-2 Spike binding to ACE2 can induce formation of multicellular syncytia (Sanders et al., 2021; Xu et al., 2020) and we used a real-time assay to assess the effect of SR9009 on Spike-dependent cell-cell fusion (Thakur et al., 2020). Initial experiments evaluated Calu-3 cells as targets in this fusion assay but due to their preferred growth in clusters we obtained high inter- and intra-assay variability. In contrast Huh-7 hepatoma cells, previously used to study SARS-CoV-1 infection (Gillim-Ross et al., 2004; Lambert et al., 2005; Nie et al., 2004), that express endogenous ACE2 and grow as a monolayer supported Spike cell-cell fusion. We confirmed that treating Huh-7 cells with SR9009 reduced ACE2 expression (**Figure 2F**) and Spike driven cell-cell fusion (**Figure 2G**). In summary, these studies demonstrate that the REV-ERB agonist SR9009 represses ACE2 expression and limits SARS-CoV-2 entry and cell-cell fusion.

BMAL1 and REV-ERB regulate gene expression by binding E-box or ROR response elements (RORE), respectively, in the promoter and enhancer regions of their target genes (Harding and Lazar, 1993). A genome-wide CRISPR screen identified 153 host factors with a potential role in SARS-CoV-2 infection (Daniloski et al., 2020) and bio-informatic analysis identified 144 canonical E-box motifs ‘CANNTG’ and 80 ROR response elements ‘RGGTCA’ in the promoter regions of these genes (**Figure 3A**), suggesting a role for circadian pathways in SARS-CoV-2 RNA replication. To evaluate a role for BMAL1 in post-entry steps in the virus life cycle we assessed SARS-CoV-2 (Victoria 01/20 strain) replication in *Bmal1* silenced Calu-3 and parental cells and observed a significant reduction in both intracellular and extracellular viral RNA along with infectious virus (**Figure 3B**). Importantly, we observed reduced replication of the Alpha (B.1.1.7 strain) and Beta (B.1.351 strain) variants of concern in the *Bmal1* silenced cells (**Figure 3C**). We next assessed whether pharmacological inhibition of BMAL1 could restrict viral replication. Treating Calu-3 cells with SR9009 or the cryptochrome stabilizer KL001 (Hirota et al., 2012) resulted in a dose-dependent reduction in SARS-CoV-2 RNA levels (**Figure 3D**). We confirm that SR9009 or KL001 treatment had no detectable effect on the Calu-3 cell cycle status under the conditions used for the infection experiments (**Figure S1B**). Importantly, both drugs significantly inhibited infection of ALI cultures of proximal airway epithelial cells (**Figure 3E**). Finally, we demonstrate that SR9009 treatment of Calu-3 cells inhibited the replication of both Alpha and Beta B.1.1.7 and B.1.351 SARS-CoV-2 variants (**Figure 3F**).

**Figure 3.**
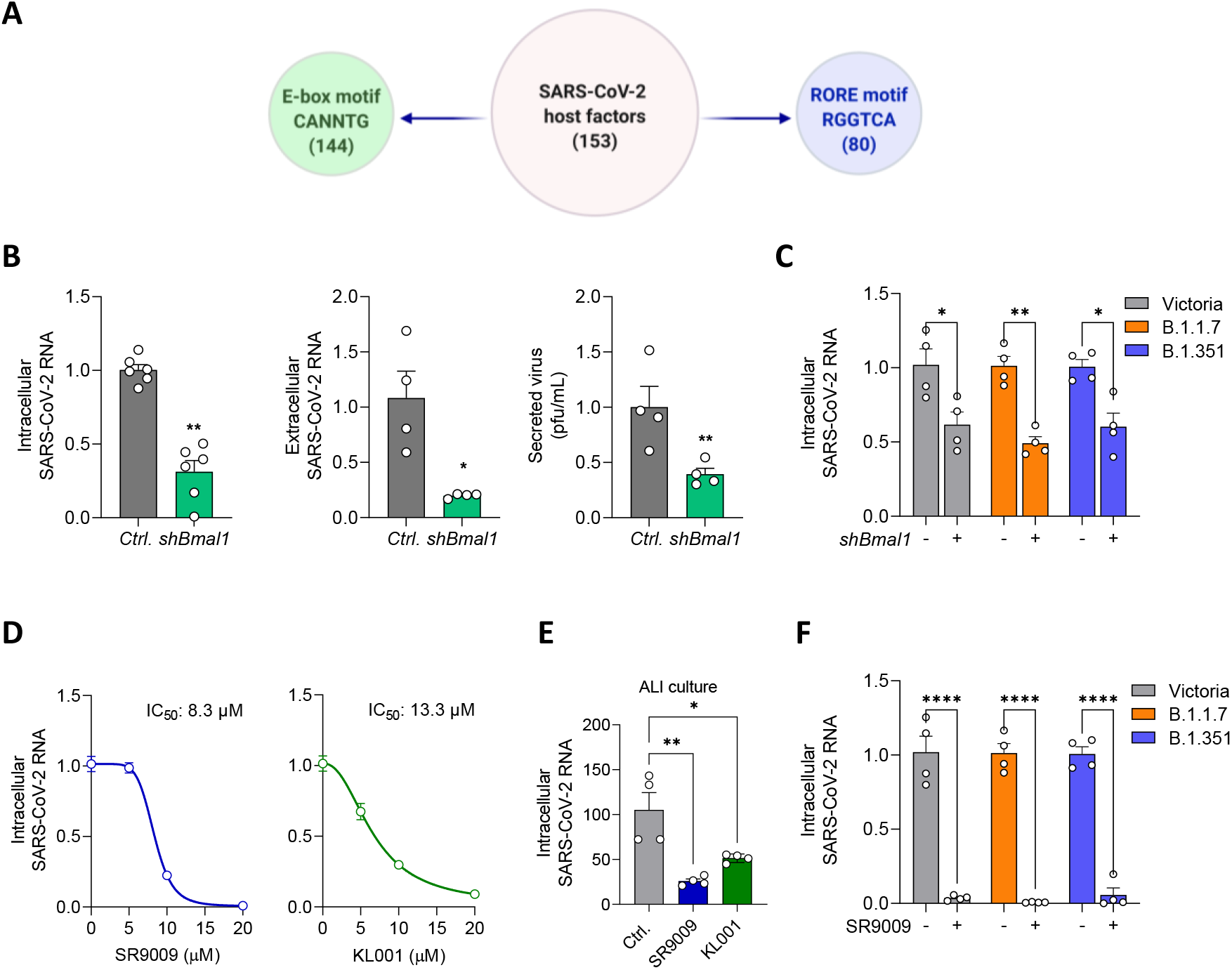
Genetic or pharmacological targeting BMAL1 reduces SARS-CoV-2 replication. (**A**) Circadian regulatory motifs in the promoter of proposed SARS-CoV-2 host factors. Sequence analysis of promoters of genes encoding SARS-CoV-2 host genes with HOMER (Hypergeometric Optimization of Motif EnRichment tool) identifies a canonical E-box motif ‘CANNTG’ in 144 of the 153 genes and an ROR response element ‘RCGTCA’ in 80. (**B**) Bmal1 silencing reduces SARS-CoV-2 replication. Calu-3 cells were transduced with lentivirus encoding shBmal1 or a control followed by SARS-Cov-2 infection. Viral RNA was measured at 24h post-infection (hpi) and expressed relative to control cells. Infectivity of the extracellular viral particles was assessed by plaque assay using Vero-TMPRSS2 cells (mean ± S.E.M., n = 4-6, Mann–Whitney test). **(C)** Calu-3 cells stably transduced with lentivirus encoding shBmal1 or a control were infected with SARS-CoV-2 Victoria 01/20, B1.1.7, or B1.351 (MOI 0.003) and viral replication assessed 24hpi by measuring intracellular viral RNA and data expressed relative to the control (mean ± S.E.M., n = 4, Mann–Whitney test). **(D)** Calu-3 cells were treated with REV-ERB agonist SR9009 or the cryptochrome stabilizer KL001 for 24h and infected with SARS-Cov-2 infection. Intracellular viral RNA was measured at 24hpi and expressed relative to the untreated control. Data are presented as mean ±S.E.M. from n = 4 independent biological replicates. **(E)** Differentiated air-liquid interface cultures of proximal airway epithelial cells were treated with SR9009 (20 μM) or KL001 (20 μM) for 24h followed by SARS-Cov2 infection. Intracellular viral RNA was quantified at 24hpi and expressed relative to the untreated control. Data are presented as mean ± S.E.M., n = 4, Kruskal–Wallis ANOVA with Dunn’s test. **(F)** Calu-3 cells were treated with SR9009 for 24h followed by infection with SARS-CoV-2 Victoria 01/20, B1.1.7, or B1.351 (MOI 0.003) and viral replication assessed at 24hpi by measuring intracellular viral RNA and data expressed relative to the untreated control (mean ± S.E.M., n = 4, Mann–Whitney test). See also Figure S1.

To define whether circadian pathways regulate post-entry steps in the SARS-CoV-2 life cycle, we evaluated the effect of SR9009 on viral replication when added before or after viral inoculation. The agonist reduced intracellular RNA and secretion of infectious particles under both conditions (**Figure 4A**), leading us to conclude that both SARS-CoV-2 entry and replication are circadian regulated. To explore the mechanism underlying the antiviral phenotype we sequenced *Bmal1* silenced and SR9009 treated Calu-3 cells. Differential expression analysis was performed and Gene Set Enrichment Analysis (GSEA) (Subramanian et al., 2005) showed an enrichment in previously defined circadian gene sets from the molecular signatures database (MSigDB) (Liberzon et al., 2011) (**Figure 4B, Figure S4A-B**). GSEA was performed to investigate host pathways regulated by BMAL1 or SR9009. Using the Hallmarks gene sets from MSigDB (Liberzon et al., 2015), we found that 32 of 50 gene sets were significantly enriched by Bmal1 silencing (**Figure 4C**) while 18 of 50 gene sets were significantly altered by SR9009 treatment (**Figure S4C**). Genes involved in energy metabolism including fatty acid metabolism and cholesterol homeostasis, as well as hypoxia pathways were enriched in both data sets. Interestingly, both cholesterol homeostasis and hypoxia pathways were previously linked to SARS-CoV-2 replication with pharmacological agents targeting these pathways showing antiviral activity (Saran et al., 2020; Wing et al., 2021; Wu et al., 2017).

**Figure 4.**
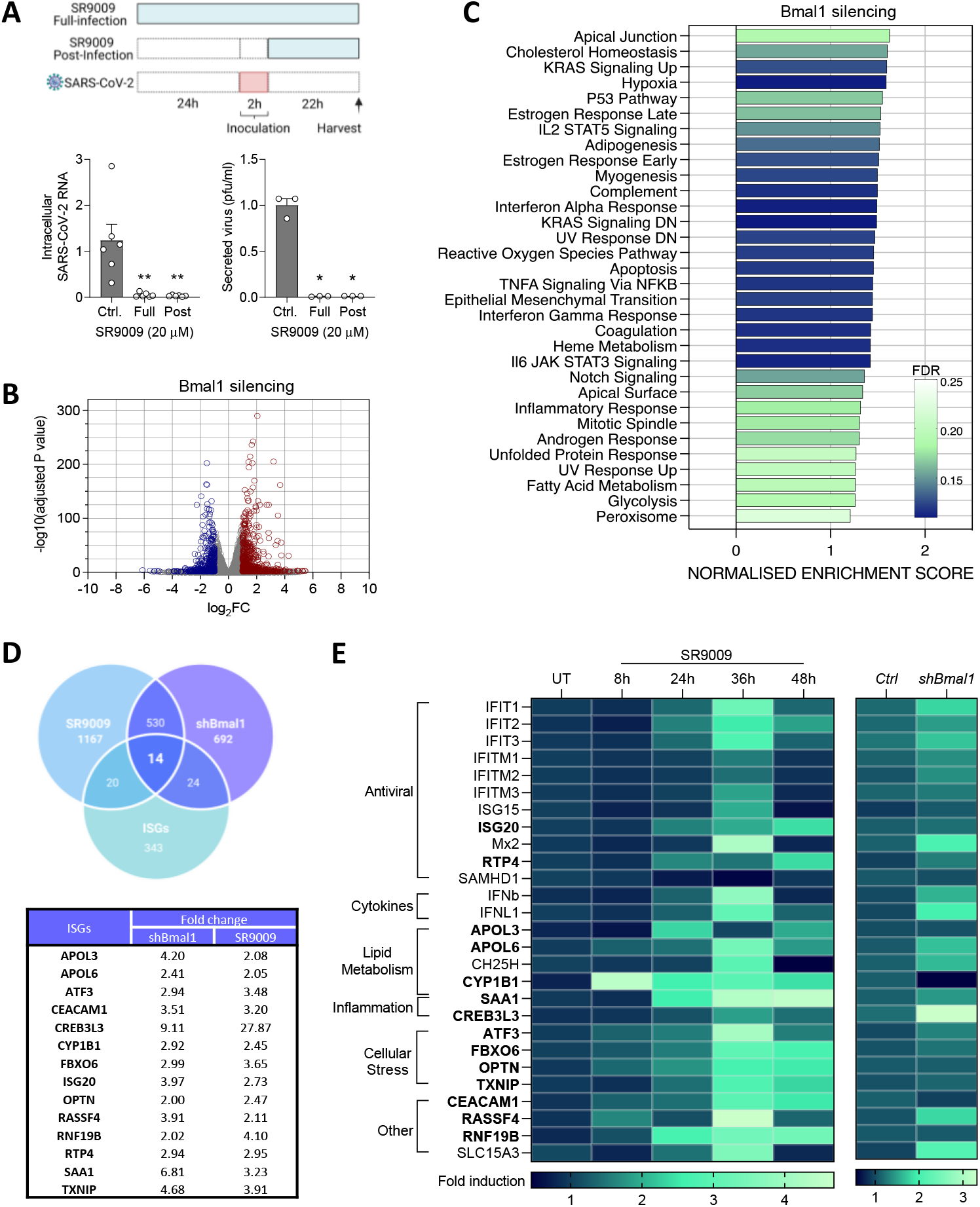
Transcriptomic analysis of Bmal1 silenced and SR9009 treated cells highlight an induction of antiviral ISG response. **(A)** Calu-3 cells were treated with SR9009 (20 µM) either 24h before infection or from 2h post-inoculation with SARS-CoV-2. Cells were incubated for 22h, intracellular viral RNA quantified by qPCR and expressed relative to the untreated control. Infectivity of the extracellular virus was assessed by plaque assay using Vero-TMPRSS2 cells. Data are presented as mean ± S.E.M. from n = 3 independent biological replicates. Statistical significance was determined using Kruskal–Wallis ANOVA with Dunn’s test. **(B)** Gene expression was quantified by RNA sequencing in Bmal1 silenced or untreated Calu-3 cells. Differential expression analysis was performed between the conditions using DESeq2 Package. Volcano plot shows significantly differentially expressed genes based on a log2FC of +/-1 and an adjusted (Benjamini Hochberg) P value of 0.05. Red points denote significant upregulation, blue denotes downregulation. **(C)** Gene set enrichment analysis (GSEA) was performed to investigate host pathways regulated by Bmal1 silencing. Using the Hallmarks gene sets from the molecular signatures database, 32 out of 50 gene sets were significantly upregulated in shBmal1 above control cells, at an FDR of less than 25%. Significantly enriched hallmarks were plotted, ranked by normalised enrichment score, and coloured by FDR. **(D)** Overlap of differentially expressed genes from Bmal1 silenced and SR9009 treated Calu-3 cells with reported Interferon-Stimulated Genes (ISGs) (Kane et al., 2016). **(E)** ISG RNA levels of Calu-3 cells treated with SR9009 for 8, 24, 36 or 48h an shBmal1 Calu-3 cells were measured by qRT-PCR and normalized to TBP (TATA box binding protein, n = 3 biological replicates). The heatmaps display the fold induction relative to the untreated control. See also Figures S4.

Interferons (IFNs) constitute the first line of defense against viral infections by inducing the expression of various interferon-stimulated genes (ISGs). Emerging evidence highlights the importance of IFNs in controlling SARS-CoV-2 infection and disease outcome (Arunachalam et al., 2020; Hadjadj et al., 2020; Zhang et al., 2020). We noted that IFN response pathways were significantly enriched in the Bmal1 silenced Calu-3 cells (**Figure 4C**). Furthermore, a recent study reported the circadian regulation of ISGs in murine skin (Greenberg et al., 2020), prompted us to investigate a link between BMAL1 and ISG expression in Calu-3 cells. We identified an overlap of 19 ISGs co-regulated by Bmal1 and SR9009 treatment with previously published ISGs (**Figure 4D**) (Kane et al., 2016).

To understand the kinetics of the induction of these ISGs, Calu-3 cells were treated with SR9009 for 8h, 24h, 36h and 48h followed by qPCR of the shBmal1/SR9009 co-regulated ISGs and additional genes either induced by SARS-CoV-2 infection or reported to show antiviral activity against SARS-CoV-2 (Martin-Sancho et al., 2021; Wang et al., 2020b). This panel showed a strong induction for most ISGs >36h post-treatment (**Figure 4E**), consistent with the antiviral activity of SR9009 at 48h in the above experimental settings. We confirmed ISG induction in Calu-3 shBmal1 cells and this was not limited to ISGs with direct antiviral activity but includes those involved in many cellular pathways (**Figure 4E**).

Our studies show evidence for circadian regulation of ACE2 in Calu-3 lung epithelial cells and show striking inhibitory effects of BMAL1 silencing, circadian modifying agents SR9009 and KL001 on SARS-CoV-2 entry and virus replication. Of note, SARS-CoV-1 and alpha NL63 also require ACE2 to enter cells (Li, 2015) and our data support a role for circadian factors regulating the infection of these related coronaviruses. The human *Ace2* promoter encodes putative binding sites for BMAL1/CLOCK and REV-ERB, however, we did not observe any binding of these factors by chromatin immunoprecipitation-qPCR in Calu-3 cells (**Figure S5**). In addition, there was no evidence for rhythmic expression of *Ace2* transcripts in Calu-3 or in the lung or liver from entrained mice or in published diurnal/circadian datasets from baboons (Mure et al., 2018) or mice (Zhang et al., 2019). These results are consistent with studies showing a role for post-transcriptional and post-translational mechanisms in circadian regulation (Kojima et al., 2011; Reddy et al., 2006). There is an emerging role for microRNAs (miRNAs) in modulating the circadian clock (Park et al., 2020; Pegoraro and Tauber, 2008) and miR-18 has been linked to ACE2 expression (Widiasta et al., 2020), providing a possible post-transcriptional mechanism. Circadian pathways are recognised to affect the pharmacokinetics and pharmacodynamics of drug responses (Ruben et al., 2019). Our study showing that ACE2 is a rhythmically ‘moving’ target could be relevant to the evaluation of new treatments targeting this step in the viral life cycle.

In addition to the circadian regulation of ACE2-mediated viral entry, we observed a marked suppression of SARS-CoV-2 RNA and genesis of infectious particles following SR9009 treatment and in Bmal1 silenced cells. Our bio-informatic analysis suggests that 30% of potential SARS-CoV-2 host factors are potentially BMAL1/REV-ERB regulated, highlighting a role for circadian-signalling to influence multiple steps in the virus life cycle. Further work is needed to characterise the circadian-dependent mechanisms of SARS-CoV-2 repression. However, our data suggest a role for BMAL1 to regulate ISGs which could impact SARS-CoV-2 replication. Namely, we can show that ISG20 and members of the IFIT family that were previously reported to have antiviral activity against SARS-CoV-2 (Martin-Sancho et al., 2021) are induced in Bmal1 silenced cells. In keeping with literature for other circadian regulated inflammatory immune responses this increase in ISG transcripts could be regulated at different stages of the IFN sensing and signalling pathway, via direct or indirect mechanisms. For example, some Toll Like Receptors (TLR) are circadian regulated, with TLR9 having non-canonical E-box motifs (Silver et al., 2012). The interplay between IFNs and COVID-19 disease progression is complex and despite initial findings there is currently no clear evidence for a genetic immune predisposition to severe COVID-19 (Meisel et al., 2021; Povysil et al., 2021). Nevertheless, there is evidence at early stages of infection that SARS-CoV-2 is sensitive to ISG induction and IFNs are being discussed as a treatment option (Cheemarla et al., 2021; Hasselbalch et al., 2021; Yang et al., 2021). In this light our findings could inform future therapeutic approaches.

A key finding from our study is the potential application of chronomodifying drugs for the treatment of SARS-CoV-2 infection (Sengupta et al., 2021). Dexamethasone is one of the few drugs that can reduce the severity of COVID-19 (Group et al., 2020) and is known to synchronise circadian pathways (Oster et al., 2006; Pezuk et al., 2012). Over the last decade, a number of compounds that target core clock proteins have been developed (Ercolani et al., 2015), including REV-ERB (Everett and Lazar, 2014; Wang et al., 2020a) and ROR (Hirota et al., 2012; Solt and Burris, 2012) that we previously reported could inhibit hepatitis C virus and HIV replication (Borrmann et al., 2020; Zhuang et al., 2019). A report demonstrating REV-ERB dependent and independent effects of SR9009 (Dierickx et al., 2019), suggests some additional off-target effects. We cannot exclude the possibility of additional pathways contributing to SR9009 anti-viral activity; however, our use of genetic targeting approaches confirms a role for BMAL1 in regulating SARS-CoV-2 replication. Since REV-ERBα agonists impact the host immune response by suppressing inflammatory mediators such as IL-6 (Gibbs et al., 2012) they offer a ‘two pronged’ approach to reduce viral replication and adverse host immune responses.

Epidemiological (Pan et al., 2011; Vyas et al., 2012) and molecular evidence (Kervezee et al., 2018) shows that night shift workers suffer circadian disruption and are at an increased risk of developing chronic inflammatory diseases. A recent study reported that shift work was associated with a higher likelihood of in-hospital COVID-19 positivity (Maidstone et al., 2021). Our observations raise questions as to how our *in vitro* studies translate to humans, where the time of exposure to SARS-CoV-2 may impact on the likelihood of infection, the host response, virus shedding, transmission and disease severity and are worthy of further investigation.

## ACKNOWLEDGEMENTS

The authors would like to thank colleagues for the provision of reagents: Pramila Rijal and Alain Townsend for anti-S neutralizing mAb F1-3A and ACE2-Fc (WIMM, Oxford). Daniel Watterson, Keith Chappell, Ariel Isaacs and Naphak Modhiran (University of Queensland) for the Spike D614G mutant. Willam James (University of Oxford) for access to VOC. Darren Blase and Zuzana Bencokova for facilitating virus work in NDMRB. We would like to thank Romain Martin and Nicolas Brignon for excellent technical assistance and Atish Mukherji for helpful discussions. The McKeating laboratory is funded by a Wellcome Investigator Award (IA) 200838/Z/16/Z, UK Medical Research Council (MRC) project grant MR/R022011/1 and Chinese Academy of Medical Sciences (CAMS) Innovation Fund for Medical Science (CIFMS), China (grant number: 2018-I2M-2-002). Mirjam Schilling is funded by the Deutsche Forschungsgemeinschaft (DFG, German Research Foundation) (SCHI 1487/2-1). The Hinks laboratory is funded by grants from the Wellcome (104553/z/14/z, 211050/Z/18/z) and the National Institute for Health Research (NIHR) Oxford Biomedical Research Centre; the views expressed are those of the authors and not those of the NHS or NIHR. Aarti Jagannath is funded by David Phillips fellowship from BBSRC-UKRI (BB/N01992X/1). The Baumert lab is funded by grants from the FRM and ANR TargEnt-COVID-19, Inserm, the University of Strasbourg, the Institut Universitaire de France and the Agence Nationale de Recherche sur le Sida et les hépatites virales (ANRS), the National Institute of Allergy and Infectious Diseases of the National Institutes of Health under award number U19AI12386, ERC AdG HEPCIR, the European Union’s Horizon 2020 research and innovation programme under grant agreement No 671231, FONDATION ARC TheraHCC2.0 and Fondation ARC pour la recherche sur le cancer (grant n° IHU201901299).

## AUTHOR CONTRIBUTIONS

XZ designed and conducted experiments and co-wrote MS; ST designed and conducted experiments and co-wrote MS; FW designed and conducted experiments; PACW designed and conducted experiments; MS designed and conducted experiments and edited MS; JMH designed and conducted experiments and ran bio-informatic analysis; HB conducted experiments and analysed data; SBM provided PBECs; JC provided ALI cultures; LM provided resources; LH and CB conducted experiments; CS designed and analyzed experiments; NZ conducted experiments; CC conducted experiments; HS designed and conducted experiments; LH conducted experiments; CB conducted experiments; AA conducted experiments; SW conducted analysis; TKH provided reagents; LS provided reagents; K-YAH provided reagents; KW provided experimental data; TSCH provided reagents; AJ provided experimental data; SV experimental data; DB designed and conducted experiments and provided reagents; TFB designed and analyzed experiments, provided reagents and co-wrote MS; JAM designed the study and co-wrote the MS.

## STAR METHODS

### RESOURCE AVAILABILITY

#### Lead contact

Further information and requests for resources and reagents should be directed to and will be fulfilled by the lead contact, Jane McKeating. Materials availability: This study did not generate new unique reagents. Data and Code availability: The authors declare that all data supporting the findings of this study are available in the article along with supplementary Information file and source data. The RNA-seq data from shBmal1 silenced or SR9009 treated Calu-3 cells are deposited at NCBI (GSE176393).

### EXPERIMENTAL MODEL AND SUBJECT DETAILS

#### Animals

Mouse experiments were carried out at the Institute of Viral and Liver Disease animal facility (approval number E-67-482-7). C57BL/6J male mice were purchased from Charles River and housed in individually ventilated cages under a 12/12 dark/light cycle with a ZT0 corresponding to 7am. After two weeks of acclimatisation, nine-week-old mice were sacrificed at different time points (ZT0, ZT4, ZT8, ZT12, ZT16, ZT20; n = 5/time point). Organs were harvested after exsanguination by intracardiac puncture, frozen in liquid nitrogen and kept at -80°C until further processing.

#### Cell culture

Calu-3, Huh-7, HEK293T and Vero E6 cells were cultured in DMEM supplemented with 10% fetal bovine serum (FBS), 2mM L-glutamine, 100 U/mL penicillin and 10μg/mL streptomycin (all reagents from Life Technologies/Thermo Fisher). Calu-3 was a kind gift from Dr Anderson Ryan (Oncology Department, University of Oxford). Vero E6/TMPRSS2 cells (a Vero E6 cell line stably overexpressing the TMPRSS2 gene, kindly provided by Dr Makoto Takeda at Department of Virology III, National Institute of Infectious Diseases (Matsuyama et al., 2020)) were cultured in DMEM supplemented with 10% fetal bovine serum, 10 units/mL penicillin, 10 mg/mL streptomycin, 10 mM HEPES (pH 7.4), and 1 mg/mL G418 (Life Technologies, UK). All cell lines were maintained at 37°C and 5% CO2 in a standard culture incubator. For Air Liquid Interface (ALI) culture, primary bronchial epithelial cells were seeded onto 0.4 µm transwells (Greiner Bio-One, Frickenhausen, Germany) and cultured in Promocell Airway Epithelial Cell Growth media until confluent. Media was removed and cells fed basally with Pneumacult-ALI media (StemCell Technologies, Vancouver, Canada) for 3 months to ensure complete differentiation to ALI with uniform ciliary beating and mucus production visible. Human primary bronchial epithelial cells were obtained using flexible fibreoptic bronchoscopy from healthy control volunteers under light sedation with fentanyl and midazolam. Participants provided written informed consent. The study was reviewed by the Oxford Research Ethics Committee B (18/SC/0361).

#### SARS-CoV-2 strains

Victoria 01/20 (BVIC01) (Caly et al., 2020) (provided by PHE Porton Down after supply from the Doherty Centre Melbourne, Australia); B.1.1.7 (Tegally et al., 2020) (20I/501Y.V1.HMPP1) (provided by PHE Porton Down) and B.1.351 (201/501.V2.HV001) (Cele et al., 2021) (Centre for the AIDS Programme of Research in South Africa) were passaged in Vero E6 cells.

### METHOD DETAILS

#### Cell synchronization and luminescence recording

Calu-3 cells were seeded at 8 x 10^4^ per cm^2^ surface area of tissue culture plastic for 24h followed by synchronisation with complete media containing 50% FBS for 1h. Serum shocked Calu-3 cells were washed twice with PBS and cultured in normal media for 12h prior to starting any time-dependent experiments as previously reported (Balsalobre et al., 1998; Pariollaud et al., 2018). For live luminescence recording, Calu-3 cells stably expressing Bmal1-luciferase promoter were seeded in 96-well plates at 2.5 x 10^4^ per well and 24h later synchronized by serum shock for 1h and incubated with Vivo-Glo (Promega, UK) and monitored in real-time at 30-min intervals for > 84h on a microplate reader (Clariostar, BMG Labtech, UK).

#### Cell cycle analysis

Calu-3 cells were incubated in 10 μM bromodeoxyuridine (BrdU) (Sigma) for 60 minutes at 37oC. Cells were then washed, and trypsinsed before fixation in ice cold ethanol. Cells underwent 30 minutes digestion in warmed pepsin solution (Sigma), before treating with 2M HCl for 15 minutes. Samples were washed, then blocked (0.5% BSA/0.5% Tween20) for 30 minutes before staining with a-BrdU-488 (Biolegend) directly conjugated antibody and Propidium Iodide (Invitrogen). Samples were acquired on BD Cyan Flow Cytometer and analysed via Flow-Jo.

#### SARS-CoV-2 pseudoparticle genesis and infection

SARS-CoV-2 lentiviral pp were generated by transfecting HEK-293T cells with p8.91 (Gag-pol), pCSFW (luciferase reporter) and a codon optimised expression construct pcDNA3.1-SARS-CoV-2-Spike, as previously reported (Thompson et al., 2020). The Furin cleavage site mutant was generated by mutagenesis of a pcDNA3.1 based clone expressing a C-terminally flag-tagged SARS-CoV-2 Spike protein (Wuhan-Hu-1 isolate; MN908947.3). The polybasic cleavage site TNSPRRA in SARS-CoV-2 Spike was replaced with the corresponding SARS-CoV variant sequence SLL. The pNBF SARS-CoV2 FL D614G mutant was a kind gift from Dr. Daniel Watterson and Dr. Naphak Modhiran at the University of Queensland. Supernatants containing viral pp were harvested at 48 and 72h post-transfection, frozen and stored at -80°C. As a control pp were generated that lacked a viral envelope glycoprotein and were included in all infection experiments to control for non-specific uptake. This control was included in all pp experiments and the luciferase values subtracted from values acquired with the SARS-CoV-2pp. To confirm spike-dependent infection, SARS-CoV-2pp were incubated with the anti-S-mAb FI-3A (1µg/mL)(Hauang et al., 2020) for 30 min prior to infection for all experiments.

SARS-CoV-2 VSV pp were generated as previously reported(Hoffmann et al., 2020b) using reagents provided by Stefan Pöhlmann (Infection biology unit, German Primate Center, Göttingen, Germany). Briefly, HEK-293T cells were transfected with an expression construct encoding a c-terminal truncated version of SARS-CoV-2-S for assembly into VSV pp (pCG1-SARS-CoV-2-S-ΔC)(Hoffmann et al., 2020a). After 24h, the transfected cells were infected with a replication-incompetent VSV (VSV*ΔG(Berger Rentsch and Zimmer, 2011)) containing GFP and firefly luciferase reporter genes, provided by Gert Zimmer (Institute of Virology and Immunology, Mittelhäusern, Switzerland). After 1h, the cells were washed with PBS before medium containing a VSV-G antibody (I1, mouse hybridoma supernatant from CRL-2700, ATCC) and supernatants harvested after 24h. The VSV*ΔG used for generating the pps was propagated in BHK-21 G43 cells stably expressing VSV-G. Viral titres were determined by infecting Calu-3 cells and measuring cellular luciferase after 48h.

#### SARS-CoV-2 propagation and infection

For infection experiments, the SARS-CoV-2 (Victoria 01/20 isolate) obtained from Public Health England was propagated in Vero E6 cells. Naïve Vero E6 cells were infected with SARS-CoV-2 at an MOI of 0.003 and incubated for 48-72h until visible cytopathic effects were observed. Culture supernatants were harvested, clarified by centrifugation to remove residual cell debris and stored at -80°C before measuring the infectious titre by plaque assay. Briefly, Vero E6 cells were inoculated with serial dilutions of SARS-CoV-2 stocks for 2h followed by addition of a semi-solid overlay consisting of 3% carboxymethyl cellulose (SIGMA). Cells were incubated for 72h, plaques enumerated by fixing cells using amido black stain and plaque-forming units (PFU) per mL calculated. For infection of Calu-3 cells, cells were plated 24h before infecting with the stated MOI. Cells were inoculated with virus for 2h after which the unbound virus was removed by washing three times with with phosphate buffered saline (PBS). Unless otherwise stated, infected cells were maintained for 24h before harvesting for downstream applications.

#### SARS-CoV-2 cell-cell fusion assay

The SARS-CoV-2 cell-cell fusion assay was performed as described previously (Thakur et al., 2020). Briefly, HEK-293T Lenti rLuc-GFP 1-7 (effector cells) and Huh-7.5 Lenti rLuc-GFP 8-11 (target cells) cells were seeded separately at 7.5×10^5^ per well in a 6 well dish in 3mL of phenol-red free DMEM, supplemented with 10% FBS, 1% sodium pyruvate and 1% penicillin/streptomycin (10,000 U/mL) before culturing overnight at 37°C, 5% CO2. The effector cells were transfected with a plasmid expressing SARS-CoV-2 Spike or a blank vector. Meanwhile, target cells were diluted to 2×10^5^/mL and 100µL seeded into a clear, flat-bottomed 96 well plate and mock-treated or treated with SR9009 (3µM or 7µM). The following day, the effector cells were harvested, diluted to 2×10^5^/mL and 100µL cultured with the drug/target cell mix, with SR9009 concentrations maintained at 3µM or 7µM. To quantify GFP expression, cells were imaged every 4h using an IncuCyte S3 live cell imaging system (Essen BioScience). Five fields of view were obtained per well at 10x magnification and GFP expression determined by calculating the total GFP area using the IncuCyte analysis software.

#### Oligonucleotides

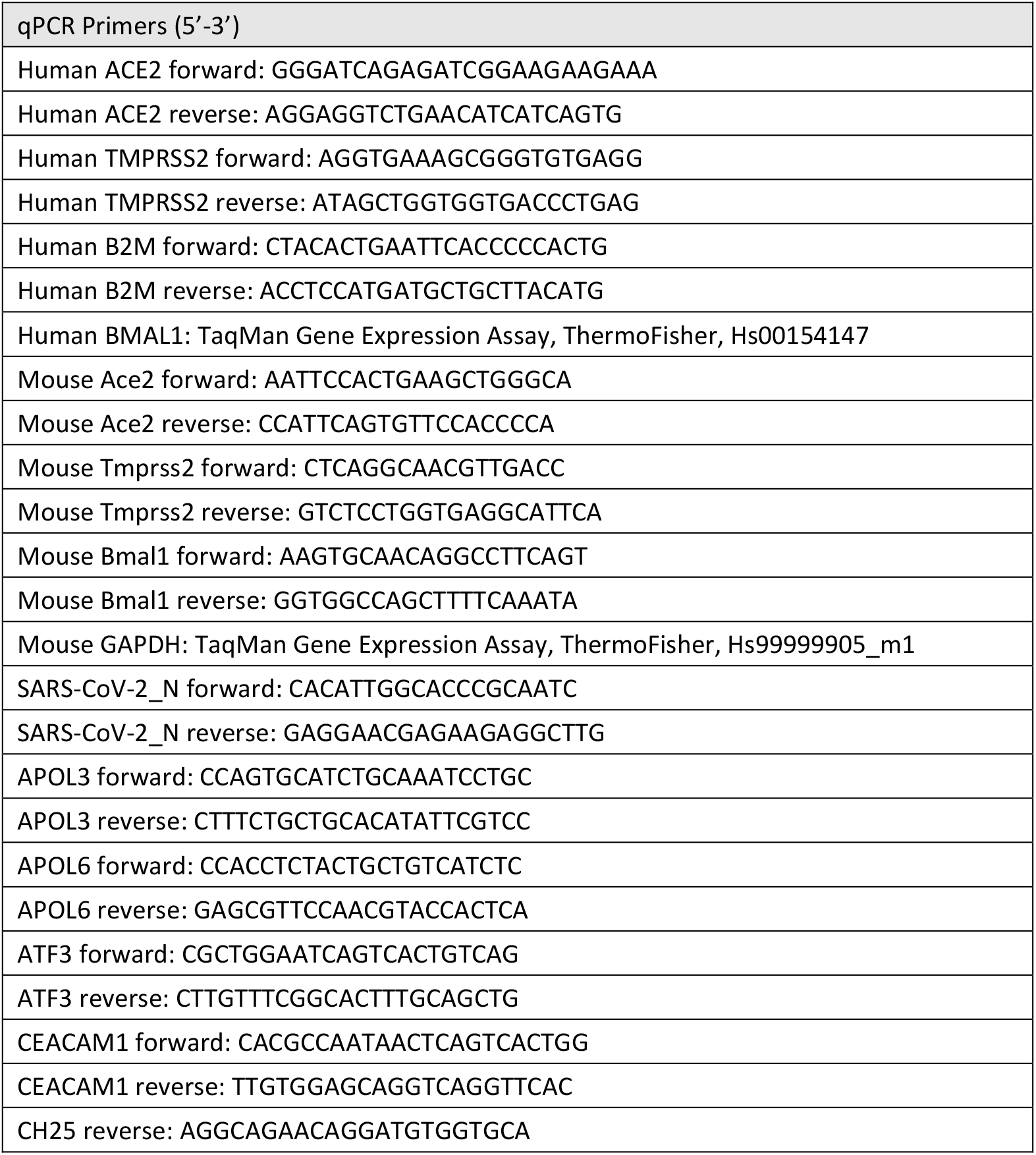
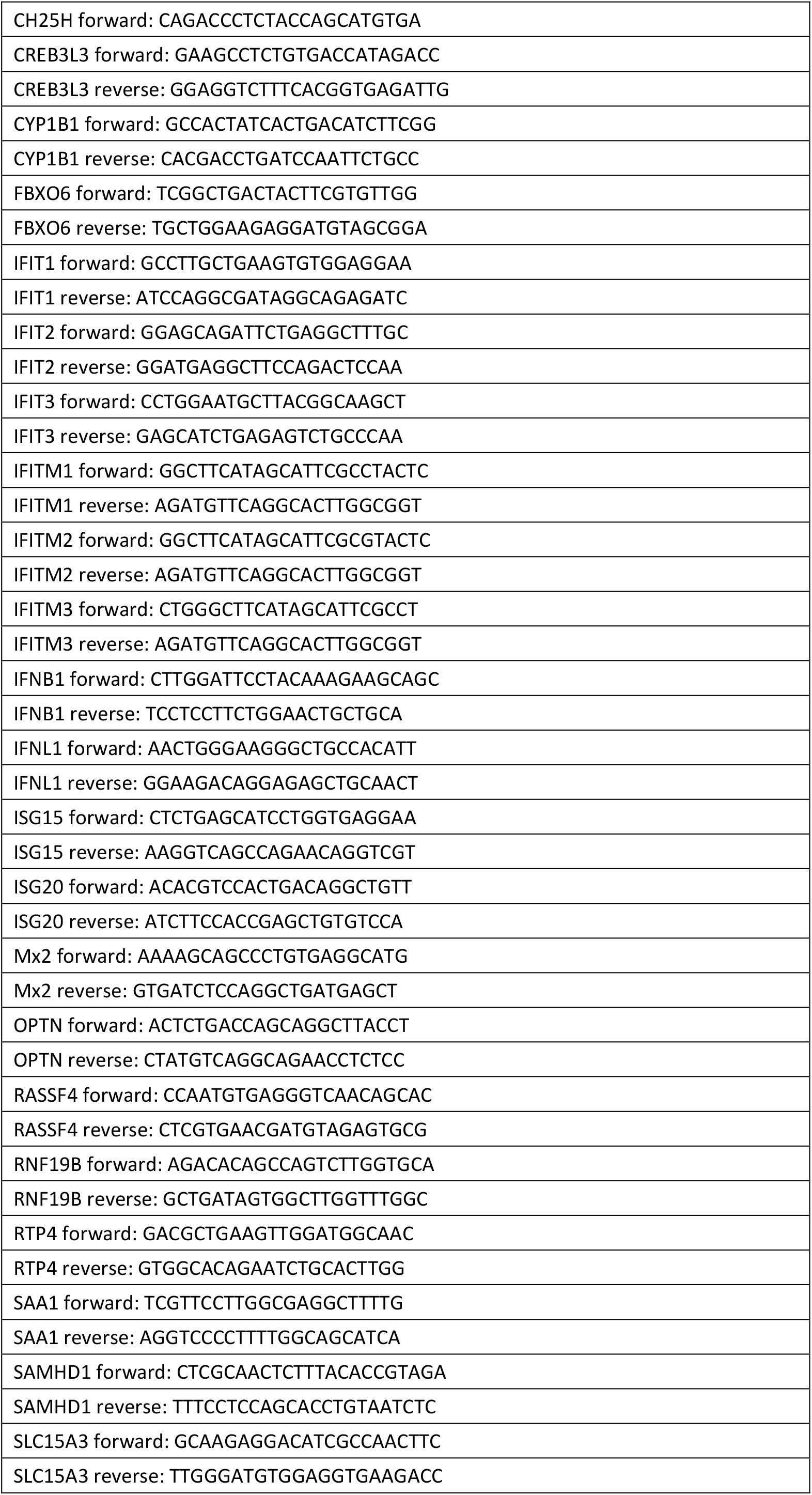
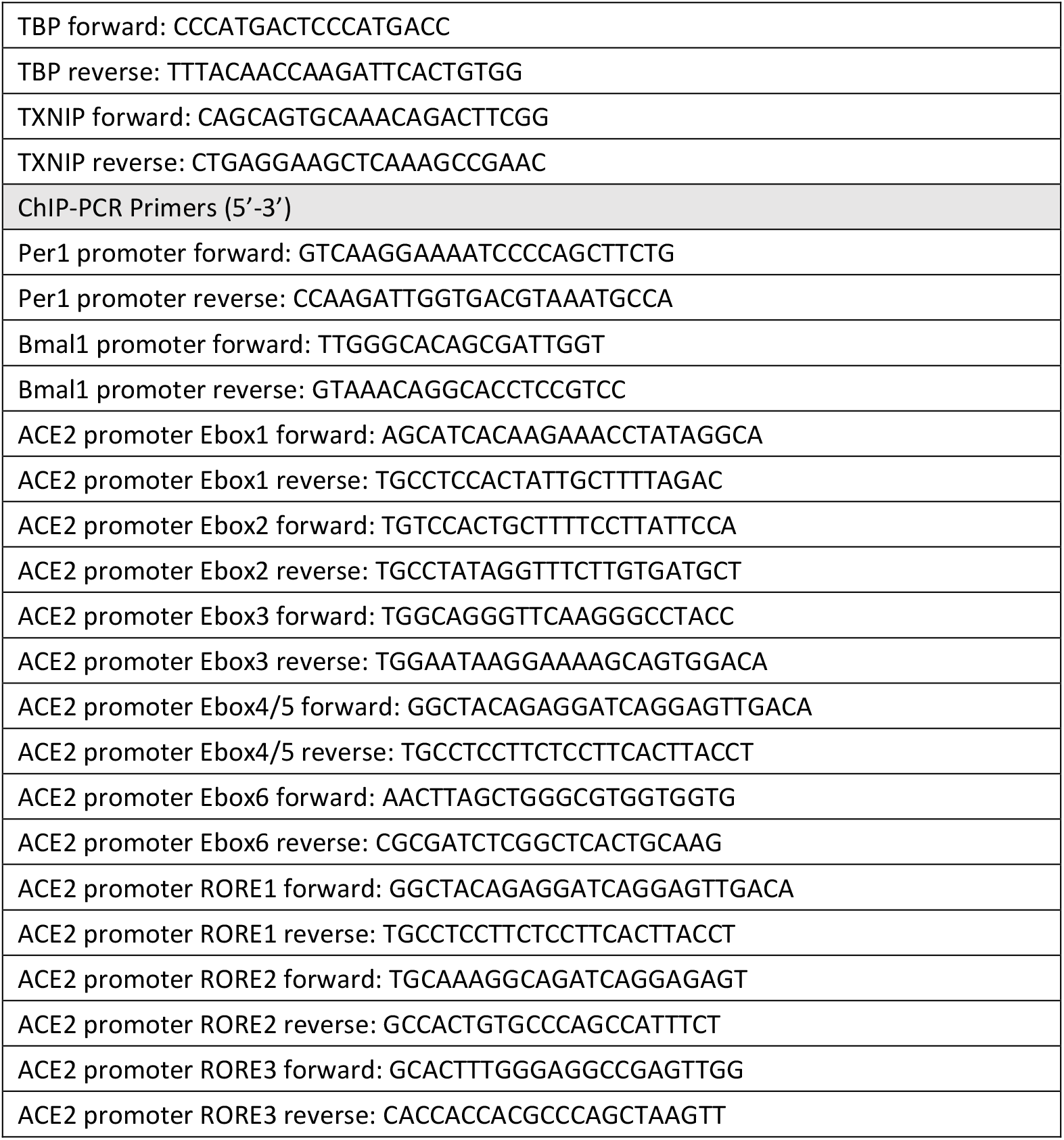

#### RT-qPCR

Cells were washed in PBS then lysed using Tri-reagent (Sigma), and mRNA extracted by phase separation or RNeasy kit (Qiagen). Equal amounts of cDNA were synthesised using the High Capacity cDNA Kit (Applied Biosystems) and mRNA expression determined using Fast SYBR master mix in a StepOne thermocycler (Applied Biosystems) using the ΔΔCt method. The lung tissues were lysed in TRI Reagent (Sigma) using a gentleMACS dissociator and RNA extracted using the manufacturers protocol. cDNAs were synthesized using the Maxima H Minus cDNA Synthesis Kit (Thermo Scientific) and mRNA quantified using iTaq Universal SYBR Green Supermix (Biorad) and specific primers on a QuantStudio 3 RT-PCR system (Applied Biosystems).

#### Immunoblotting

Cell lysates were prepared by washing cells with PBS, then lysed in Igepal lysis buffer (10mM Tris pH 7.5, 0.25M NaCl, 0.5% Igepal) supplemented with protease inhibitor cocktail (Roche Complete™) at 4°C for 5 min, followed by clarification by centrifugation (3 min, 12,000 rpm). Supernatant was mixed with Laemmli sample buffer, separated by SDS-PAGE and proteins transferred to polyvinylidene difluoride membrane (Immobilon-P, Millipore). Membranes were blocked in 5% milk in PBS/0.1% Tween-20, then incubated with anti-ACE2 (Abcam ab108252), anti-TMPRSS2 (SCBT sc-515727), anti-BMAL1 (Abcam Ab93806) or Anti-β-actin (Sigma A5441) primary antibodies and appropriate HRP-conjugated secondary antibodies (DAKO). Chemiluminescence substrate (West Dura, 34076, Thermo Fisher Scientific) was used to visualize proteins using a ChemiDoc XRS+ imaging system (BioRad). Anti-β-actin-HRP conjugate (Abcam ab49900) and/or Coomassie brilliant blue staining was used to verify equal protein loading and densitometric analysis performed using ImageJ software (NIH).

#### Bioinformatics

The published 153 SARS-CoV-2 host factors were converted to Entrez gene names (Daniloski et al., 2020). BMAL1 regulated genes were obtained from the published BMAL1 ChIP-seq in the mouse liver (Beytebiere et al., 2019). REV-ERB regulated genes were defined from published liver-specific loss of the REV-ERB paralogues (Cho et al., 2012). Promoters (−1kb from TSS) of genes encoding SARS-CoV-2 host factors were analyzed with the HOMER (Hypergeometric Optimization of Motif EnRichment) tool for motif discovery (E-box motif CANNTG; RORE motif RGGTCA). Gene set enrichment analysis was performed using GSEA_4.1.0, and gene sets of interest retrieved from the Molecular Signatures Database. Analysis was carried out using Phenotype grouping between shBmal1 and untreated Calu-3 cells, with 1000 permutations.

#### RNA-seq analysis

Bmal1 silenced, SR9009 treated and control Calu-3 cells were harvested for RNA isolation and RNA-sequencing at Novogene. RNA purity was assessed with a NanoDrop 2000 spectrophotometer (Thermo Fisher Scientific) and integrity determined using a 2100 Bioanalyzer Instrument (Agilent Technologies). Sequence adapters were removed and reads trimmed by Trim Galore v0.5.079. The reads were mapped against the reference reference human genome (hg38/GRCm38) using STAR v2.5.380. Counts per gene were calculated using Rsubread v1.28.181. Reads were analyzed by edgeR v3.30.082, normalized using TMM, counts per million calculated and differential expression analysis performed.

#### Chromatin immuno-Precipitation (ChIP) and quantitative PCR

1×10^7^ Calu-3 cells were harvested from 80% confluent 15cm plates and fixed with 1% formaldehyde (Sigma Aldrich 47608) for 10 min at room temperature before quenching with 125 mM glycine. Cells were washed twice with ice cold PBS, pelleted (800rpm, 10 min 4°C) and lysed in 500µl of Nuclear Extraction buffer (10 mM Tris pH 8.0, 10 mM NaCl, 1% NP-40) supplemented with a protease inhibitor cocktail (Roche Complete™). Samples were diluted 1:1 in ChIP Dilution Buffer (0.01% SDS, 1.1% Triton, 0.2mM EDTA; 16.7 mM Tris pH8.1, 167mM NaCl) and pulse sonicated using a Bioruptor R sonicator (Diagenode, U.K.) at high power for 15 min at 4°C (15 sec on, 15 sec off). Sonicated lysates were clarified by centrifugation at 1300 rpm for 10 min and precleared with Protein A agarose beads (Millipore, 16-156). Samples were immunoprecipitated with primary anti-BMAL1 (ChIP grade, Abcam Ab3350) or anti-REV-ERBα (ChIP grade, Abcam Ab181604) or IgG control mAb and precipitated with Protein A agarose beads. Precipitates were washed in low salt buffer (0.1% SDS, 1% Triton, 2 mM EDTA, 20 mM Tris pH8.1, 150mM NaCl), high salt buffer (0.1% SDS, 1% Triton, 2mM EDTA, 20 mM Tris pH 8.1, 500 mM NaCl), LiCl Buffer (1% Igepal, 1mM EDTA, 10 mM Tris pH 8.1, 250 mM LiCl, 1% sodium deoxycholate) and finally twice in TE wash buffer (10 mM Tris pH8.0, 1mM EDTA) before being eluted from the beads in 240µL of elution buffer (0.1 M NaHCO3, 1% SDS). Complexes were reverse crosslinked in a heated shaker at 65°C overnight, 1400 rpm, in the presence of 200 mM NaCl. Eluates were treated with Proteinase K (SIGMA) and RNaseA (SIGMA) before cleanup using MiniElute PCR Purification columns (Qiagen). Samples were analysed on a LightCycler96 (Roche) using SYBR green qPCR mastermix (PCR Biosystems, UK). Fold enrichment was calculated for each sample relative to their own IgG controls.

#### Materials

All reagents and chemicals were obtained from Sigma-Aldrich (now Merck) unless stated otherwise. REV-ERB agonist SR9009 was purchased from Calbiochem, US, dissolved in dimethyl sulfoxide (DMSO) and cytotoxicity determined by a Lactate dehydrogenase (LDH) assay (Promega, UK) or MTT assay (Sigma, UK). The BMAL1 promoter was amplified from genomic DNA using forward primer: 5ʹ-CCGCTCGAGGGGACAACGGCGAGCTCGCAG-3ʹ with reverse primer: 5ʹ-CCCAAGCTTCGGCGGCGGCGGCGGCAAGTC-3ʹ and cloned into the pGL3 luciferase reporter vector (Promega, UK). The lenti-shBmal1 plasmid was purchased from Abmgood (UK). Recombinant ACE2-Fc was previously reported(Zhou et al., 2020). The BMAL1 promoter luciferase reporter vector was purchased from Addgene, UK (Plasmid #46824) and contains 1 kb of Bmal1 upstream region and 53 nucleotides of exon 1, fused in-frame to the luciferase coding region followed by 1 kb of Bmal1 3ʹUTR. This construct has been widely used in circadian research (Brown et al., 2005; Bu et al., 2018; Sanghani et al., 2020; Schmitt et al., 2018). The Cry1 promoter construct and Bmal1/Clock expression plasmids were a gift of Ximing Qin, Anhui University, Hefei, China. The Cry1 promoter was amplified from genomic DNA using forward primer: 5ʹ-ATCCTCGAGGTAAAGATGCACATGTGGCCCTG-3ʹ and reverse primer: 5ʹ-CTAAAGCTTCGTCCGGAGGACACGCATACC-3ʹ and cloned into the pGL3 luciferase reporter vector (Promega, UK). The Bmal1 expression plasmid was previously described (Yang et al., 2020) and further engineered with a Flag tag. The Clock expression plasmid was previously described (Schoenhard et al., 2003).

### QUANTIFICATION AND STATISTICAL ANALYSIS

Data was analysed using GraphPad Prism version 8.0.2 (GraphPad, San Diego, CA, (USA). P values < 0.05 were considered significant; significance values are indicated as *p < 0.05; **p < 0.01; ***p < 0.001; ****p < 0.0001. Please see individual figure legends for further details. Curve fit for circadian synchronized cells were generated using the non-linear regression (curve-fit with sine wave) function using Graph Pad Prism 7 (GraphPad, USA).

