## SUPPLEMENTARY INFORMATION for "The circadian clock component BMAL1 regulates SARS-CoV-2 entry and replication in lung epithelial cells"

### Circadian regulation of SARS-CoV-2 infection in lung epithelial cells

Shared first authorship*

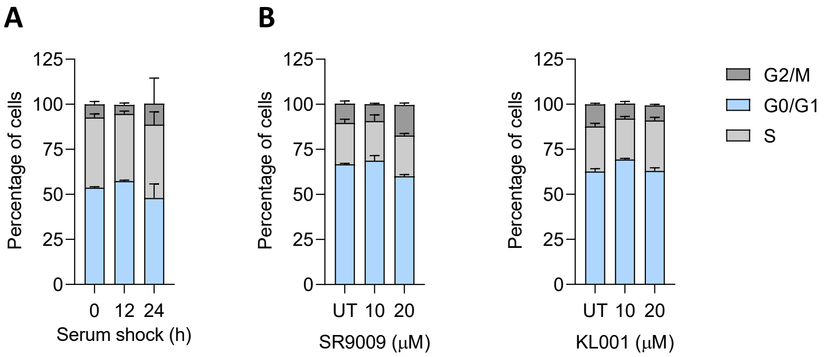


**Figure S1 Cell cycle analysis of Calu-3 cells.**

(**A**) Calu-3 cells were seeded at 8 x $10^5$/well of a 6-well plate and cell cycle analysis performed at 12h and 24h post-serum shock or (**B**) 48h post SR9009 or KL001 treatment. The proportion of cells per phase was determined by detecting BrdU incorporation and Propidium Iodide staining by flow cytometry. Statistics were tested with Two-way ANOVA (n = 3 per time point) and revealed no significant differences in cell cycle phase across time points**. (Related to Fig.1 and Fig.3)**

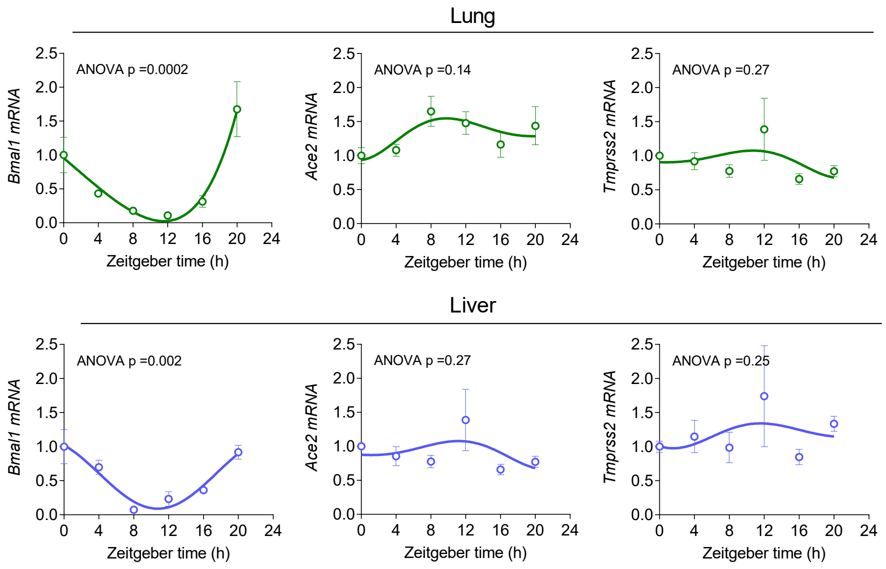


**Figure S2.** ***Bmal1, Ace2 and Tmprss2*** **transcripts in mouse tissues.**
Lung and liver tissues from circadian entrained mouse were harvested at 4h intervals and *Bmal1, Ace2* and *Tmprss2* mRNAs quantified by qPCR and expressed relative to CT0. Data represent the mean ± S.E.M., n = 5, Kruskal–Wallis ANOVA with Dunn's test. (**Related to Fig.1**).

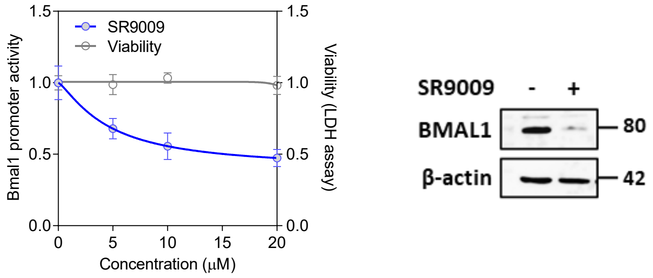


**Figure S3. Effect of REV-ERB agonist SR9009 on BMAL promoter activity and expression.**
Calu-3 cells stably expressing a Bmal1 promoter driven luciferase reporter were treated with SR9009 for 24h and promoter activity quantified by measuring luciferase activity. Cytotoxicity was determined using an LDH assay. Data are expressed relative to untreated cells and represent the mean$\pm$S.E.M., n = 16. Calu-3 cells were treated with SR9009 (20 μM) for 24h and assessed for BMAL1 expression together with housekeeping β-actin by western blotting. **(Related to Fig.2).**

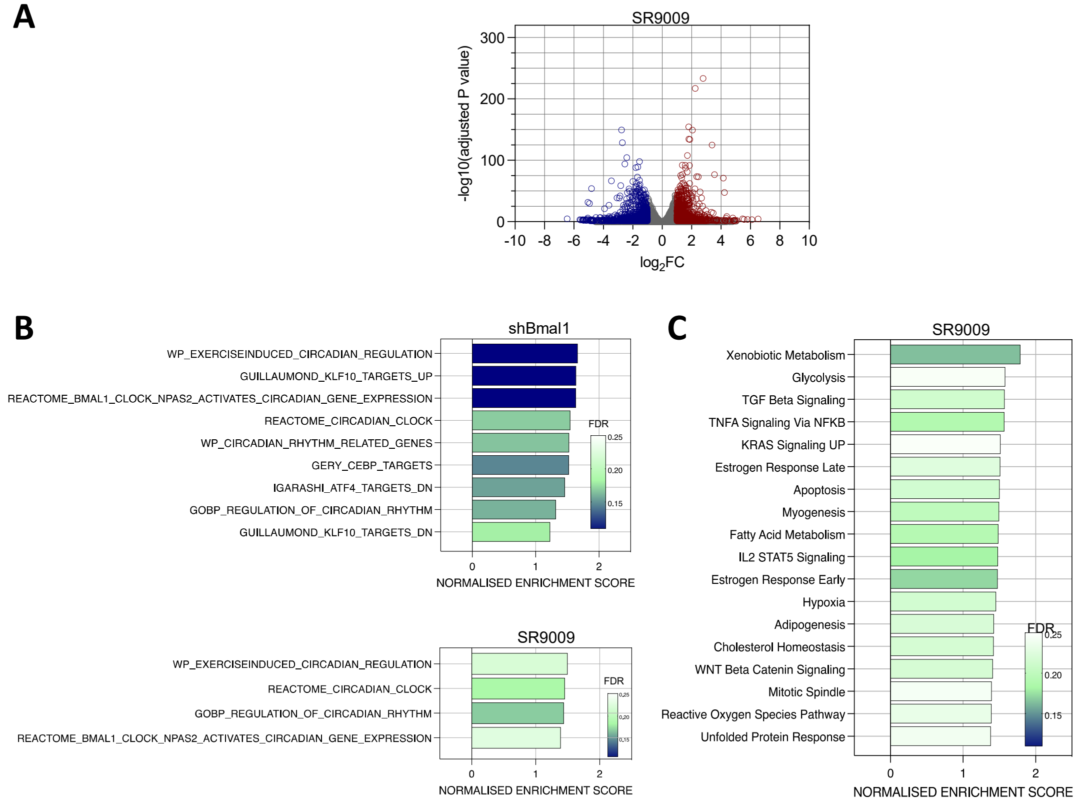


**Figure S4. RNA-seq analysis of Bmal1 silenced and SR9009 treated Calu-3 cells.**

(**A**) Differentially expressed genes in SR9009 treated Calu3 cells. Calu-3 cells were treated with SR9009 (20 µM) for 24h and gene expression quantified by RNA sequencing. Differential expression analysis was performed using DESeq2 Package between SR9009 and untreated cells. Volcano plot shows significantly differentially expressed genes based on a log2FC of +/- 1 and an adjusted (Benjamini Hochberg) P value of 0.05. Red points denote significant upregulation, blue denotes downregulation. (**B**) Validation of RNA-seq data on previously reported clock genes. All Gene sets from MSigDB were searched for the term "Circadian", and 24 gene sets obtained. GSEA was performed between shBmal1 and SR9009 against untreated cells. Gene sets were filtered for size based on >15 and <500 genes, and for their expression in both experimental groups. 9/12 circadian gene sets were significantly enriched in shBmal1, and 4/13 SR9009 treated cells. Gene sets are ranked by NES and coloured by FDR. (**C**) Gene set enrichment analysis (GSEA) for SR9009 regulated host pathways. Using the Hallmarks gene sets from the molecular signatures database, 18 out of 50 gene sets were significantly upregulated in SR9009 above untreated cells, at an FDR of less than 25%. Significantly enriched hallmarks were plotted, ranked by normalised enrichment score, and coloured by FDR. (**Related to Fig.4**).

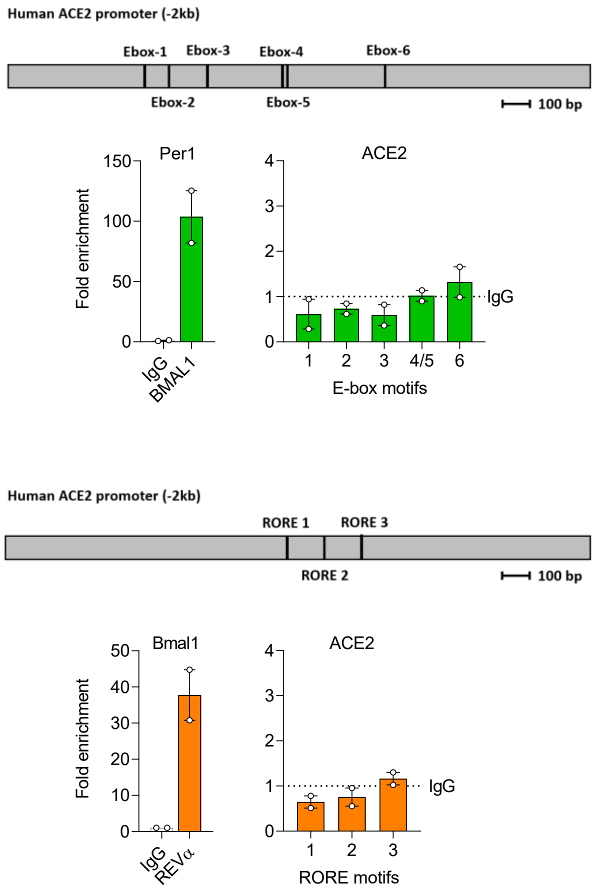


**Figure S5. ChIP analysis of BMAL1 or REV-ERBα binding ACE2 promoter in Calu-3 cells**.
Chromatin extracts from Calu-3 cells were immunoprecipitated using antibodies specific for BMAL1, REV-ERBα or rabbit IgG. qRT-PCR for ACE2 promoter DNA using primers over defined Ebox and RORE motifs or control host gene Per1 for BMAL1 ChIP and Bmal1 for REV-ERBα (positive control) was performed. IP data is presented relative to the rabbit IgG control shown as the dotted line (mean ± S.E.M., n = 2, Kruskal–Wallis ANOVA with Dunn's test).
